# Polysorbate 80 Controls Morphology, Structure and Stability of Human Insulin Amyloid-Like Spherulites

**DOI:** 10.1101/2021.08.10.455855

**Authors:** Xin Zhou, Dirk Fennema Galparsoro, Anders Østergaard Madsen, Valeria Vetri, Marco van de Weert, Hanne Mørck Nielsen, Vito Foderà

## Abstract

Amyloid protein aggregates are not only associated with neurodegenerative diseases and may also occur as unwanted by-products in protein-based therapeutics. Surfactants are often employed to stabilize protein formulations and reduce the risk of aggregation. However, surfactants alter protein-protein interactions and may thus modulate the physicochemical characteristics of any aggregates formed. Human insulin aggregation was induced at low pH in the presence of varying concentrations of the surfactant polysorbate 80. Various spectroscopic and imaging methods were used to study the aggregation kinetics, as well as structure and morphology of the formed aggregates. Molecular dynamics simulations were employed to investigate the initial interaction between the surfactant and insulin. Addition of polysorbate 80 slowed down, but did not prevent, aggregation of insulin. Amyloid spherulites formed under all conditions, with a higher content of intermolecular beta-sheets in the presence of the surfactant above its critical micelle concentration. In addition, a denser packing was observed, leading to a more stable aggregate. Molecular dynamics simulations suggested a tendency for insulin to form dimers in the presence of the surfactant, indicating a change in protein-protein interactions. It is thus shown that surfactants not only alter aggregation kinetics, but also affect physicochemical properties of any aggregates formed.

## 1. Introduction

Both *in vitro* and *in vivo* and in destabilizing conditions, proteins may undergo self-assembly processes leading to the formation of a variety of aggregates. A specific class of these aggregates, named amyloids, are implicated in the onset of human pathologies such as Alzheimer’s and Parkinson’
ss diseases (Taylor et al. 2002, Knowles et al. 2014) and type 2 diabetes (Kahn et al. 1999, Höppener and Lips 2006). Also, the impact of protein aggregates in the development of protein therapeutics may be significant, resulting in loss of active material (Wang et al. 2010, van de Weert and Randolph 2013), clogging of needles and infusion lines (van de Weert and Randolph 2013), and be a potential risk factor for unwanted immunogenicity in patients (Jiskoot et al. 2012). However, due to their exceptional strength and elasticity, self-assembled protein structures are also investigated for potential applications as biomaterials (Otzen and Nielsen 2008, Knowles and Buehler 2011, Knowles and Mezzenga 2016), for biosensing (Hauser et al. 2014), drug delivery (Maji et al. 2008), and water purification (Bolisetty and Mezzenga 2016).

Among the different protein aggregates, elongated β-sheet-rich structures named amyloid fibrils are the most studied (Dobson 1999, Dobson 2004, Gallardo et al. 2020). Moreover, both *in vitro* and *in vivo* proteins may also self-assemble into other amyloid-like forms, so-called *superstructures* (Krebs et al. 2009, Babenko et al. 2011, Foderà and Donald 2014, Vetri and Foderà 2015). Within this class, amyloid spherulites are characterized by a fascinating architecture with a spherical symmetry consisting of a central dense core from which a low-density corona grows radially (Krebs et al. 2004, Krebs et al. 2005, Vetri and Foderà 2015). Their radius spans from few µm to mm and they are readily detectable by cross-polarized light microscopy due to their specific Maltese cross pattern originating from the radial symmetry imposed by the corona (Cannon and Donald 2013). *In vitro*, and especially for globular proteins, spherulites formation can be thermally induced and often occur together with amyloid fibrils (Krebs et al. 2009, Smith et al. 2012, De Luca et al. 2020) and they are also observed *in vivo* in connection with Alzheimer’
ss disease (Exley et al. 2010, House et al. 2011).

Isolating the key interactions determining size, morphology and structural features of amyloid species is pivotal. Indeed, such a variability may reflect different biological roles (Breydo and Uversky 2015, Iadanza et al. 2018) both in neurodegenerative diseases (Taylor et al. 2002, Petkova et al. 2005, Cohen et al. 2015) and in terms of immunological responses (Bergström et al. 2005). Also, in the context of therapeutics development, the diversity of protein composites offers an opportunity for realization of tailored materials (Fennema Galparsoro et al. 2021, Norrild et al. 2021). This could result in materials e.g. for encapsulation of different active compounds (Maji et al. 2008, Shimanovich et al. 2015) and materials with tunable rheological properties (Haines-Butterick et al. 2007, Branco et al. 2009).

Co-solvents are often employed to tune protein-protein and protein-solvent interactions (PPIs and PSIs, respectively) during the aggregation process and disentangle the role of different PPIs in the reaction (Vagenende et al. 2009, Bucciarelli et al. 2020). Among the different co-solvents, surfactants are commonly used in pharmaceutical sciences to tune protein formulation stability. Surfactants interact with proteins through electrostatic and hydrophobic interactions (Otzen 2011) and reduce aggregation mainly via suppressing protein adsorption at surfaces (Parkins and Lashmar 2000); which is known to catalyze the protein refolding reaction (Randolph and Jones 2002). Moreover, surfactants may directly bind to hydrophobic patches on the protein surface, thereby reducing direct PPIs (Jones 1992, Randolph and Jones 2002, Wang et al. 2008) and lead to an overall enhanced stability of the protein ensemble (Chou et al. 2005). Notably, the direct binding between surfactant molecules and the hydrophobic domains of partially unfolded protein often occurs at the critical micelle concentration (CMC) of the surfactant (Randolph and Jones 2002, Chou et al. 2005), i.e. the bulk concentration above which the surfactant starts to self-assemble into micelles.

While the enhanced protein stability induced by surfactants is well documented, very limited information is available on the effect of surfactants on protein aggregation processes and on the features of the final aggregate structures, the latter being of vital importance for risk assessment of therapeutics. Indeed, secondary structure content, size, morphology and geometrical packing may alter the capability of the aggregates to induce immunoresponse (Filipe et al. 2010). Siposova et al. (2019) reported that the non-ionic surfactant Triton™ X-100 affected insulin amyloid aggregation at acidic pH depending on the protein:surfactant ratio, demonstrating that at concentrations below CMC insulin fibrillation was partially inhibited and fibril morphology was greatly altered. Inhibition of *in vitro* fibril formation was observed on a time scale of approximately 2 hours when the concentration of detergent was above the CMC (Siposova et al. 2019); this being ascribed to different binding phenomena between Triton™ X-100 and insulin dimers.

In the present work, we aim at investigating the mechanisms by which polysorbate 80 (PS80) affects the stability and aggregation propensity of human insulin, with a special focus on the formation of aggregates other than fibrils, i.e. amyloid spherulites, and the related structural and morphological features. PS80 is a non-ionic surfactant and one of the main excipients added in protein formulations to maintain protein therapeutic activity and prolong the shelf life of the formulations (Jones et al. 2018). Using a combination of spectroscopy and imaging methods, we reveal that PS80 does not inhibit the thermal aggregation process for insulin on a time scale of approximately 2 days, but rather delays the process. Surprisingly, addition of PS80 leads to an increase in β-sheet structure and concomitant altered morphology and enhanced stability of the formed spherulites. This demonstrates the importance of not only investigating the impact of excipient concentration on stability and aggregation kinetics, but also on the properties of the resulting aggregates, as these altered characteristics may impact safety and efficacy of the protein drug product.

## 2. Materials and Methods

### 2.1 Materials

Human insulin (91077C, ⩾95%), Thioflavin T (ThT, ≥ 65%), polysorbate 80 was obtained from Sigma Aldrich (Saint Louis, MO, USA, ∼70% oleic acid). Boric acid (99.5%-100.5%) and sodium hydroxide (≥ 99.0 %) was acquired from Merck (Darmstadt, Germany). Acetic acid was obtained from VWR Chemicals (Leuven, Belgium, ≥ 98%). Sodium hydrochloride was bought from Th. Geyer, CHEMSOLUTE (Roskilde, Denmark, 99.0%).

### 2.2 Sample Preparation

3 mg/mL (0.52 mM) insulin was dissolved in 20% (v/v) acetic acid and 0.5 M NaCl with the addition of increasing concentration of PS80. The pH value of the solution was approximately 1.7. 1 mL of sample was filtered through a 0.22 µm cellulose acetate (CA) filter (Labsolute, Th.Geyer, Renningen, Germany) into a 1.5 mL polypropylene microtube (SSIbio, Lodi, CA, USA). The concentration of insulin was determined by absorbance at 276 nm using an extinction coefficient of 1.0, measured with Nanodrop 2000C (Thermo Scientific, Wilmington, NC, USA). The concentration is presented as an average of three measurements. The aggregation was then thermally induced at 45 °C over 42 hours under quiescent conditions. Parafilm was used to seal the microtubes to limit evaporation. Residual concentration of soluble insulin not converted into aggregated after formation of the spherulites was also measured with Nanodrop 2000C (Thermo Scientific, Wilmington, NC, USA). The aggregation was then thermally induced at 45 °C over 42 hours under quiescent conditions. The formulation of insulin at acidic pH and incubation at high temperatures guarantee an *in vitro* aggregation reaction occurring in the range of several hours. This avoids extremely long aggregation reaction, which may induce artefacts in the experiments, e.g. due to sample evaporation or excessive chemical degradation. At acidic pH and in presence of acetic acid, the conditions in this study, insulin is mainly in a monomeric state (Attri et al. 2010, Schack et al. 2018), which is recognized as the building block for amyloid aggregate formation (Ahmad et al. 2003). As a consequence, such a formulation avoids the formation of higher order oligomers as dimers (occurring in presence of HCl (Attri et al. 2010)) or hexamers appearing at physiological pHs and in presence of Zn^2+^ (Attri et al. 2010), which may delay the aggregation process.

### 2.3 Thioflavin T (ThT) Fluorescence Kinetics

A plate reader system (CLARIOstar, Bmg Labtech, Ortenberg, Germany) was used to monitor the ThT fluorescence. 200 µL solution was added in each well of 96-well black polystyrene plates (Nunc, Thermo Fisher, Rochester, NY, USA), and for each sample there were four replicates. The plates were covered by polyolefin film (Thermo Fisher, Rochester, NY, USA) to avoid evaporation during the experiment. Samples were incubated at 45 °C without shaking for up to 42 hours. A stock solution of ThT consisting of dissolved and filtered (0.22 µm CA filter) ThT in purified water was stored at 4 °C until use. The final concentration of ThT was 1 mM, determined by Nanodrop at 412 nm using an extinction coefficient of 36,000 M^-1^ cm^-1^ (Groenning 2010). 20 µL of ThT stock solution was added per 1 mL insulin solutions prior to distribution in the plate with subsequent incubation. The emission intensity at 480 nm was recorded upon excitation at 450 nm every 309 s throughout the incubation time. The experiment was repeated on four different batches with four replicates for each batch (16 kinetic curves obtained for each experimental condition). Results showed comparable profiles and data trends and a representative dataset for one batch is shown in the manuscript. The time needed to reach 50% of the maximum fluorescence, t50%, was determined as a measure of the time scale of amyloid aggregation kinetics (Foderà et al. 2008).

### 2.4 Surface Tension

Surface tension of PS80 solutions was determined by Force Tensiometer K100 (Krüss, Hamburg, Germany), measuring with the rod PL03 (Krüss, Hamburg, Germany) at room temperature (RT). The CMC was roughly identified as the breakpoint in the plot of surface tension versus concentration of surfactant. PS80 was dissolved in 20 %(v/v) acetic acid with 0.5 M NaCl and data were collected at different PS80 concentrations. For each measurement, 5 mL of solution was added in the polytetrafluoroethylene (PTFE) vessel SV01 (Krüss, Hamburg, Germany). Each measurement was taken after allowing 1 min for equilibration; data were recorded until the change was smaller than 0.01 mN/m or for up to 2 min. Reported data were obtained as an average of measurements of three individual solutions with the same PS80 concentration.

### 2.5 High Performance Liquid Chromatography for Polysorbate 80 Content Estimation

The content of PS80 in the spherulites was determined by high performance liquid chromatography with evaporative light scattering detector (HPLC-ELSD). Specifically, the PS80 amount left in the supernatant after the formation of the insulin spherulites was estimated. The supernatant was obtained by centrifuging the spherulites samples at 9,279 g for 10 min and 10 °C, using Centrifuge 5417R (Eppendorf, Hamburg, Germany). As controls, PS80 solvents without insulin were incubated under the same conditions as the insulin samples (45 °C). The details of HPLC analysis regarding mobile phases, gradient and column are reported in the supplementary materials.

1 mL insulin spherulites samples after incubation were named as (Ins_PS80) here. 1 mL PS80 solution was incubated (PS80_incubated) under the same conditions (45 °C for 42 hours) as for the preparation of Ins_PS80 samples. 1 mL of PS80 solution (without insulin) was kept in the fridge (PS80_Time0) as a control to evaluate potential evaporation induced by thermal incubation at 45 °C. After 42 hours, the samples were centrifuged for 10 min at 9,279 g and 10 °C; 0.5 mL supernatant was then removed and further centrifuged 10 min at 9,279 g and 10 °C. After these steps, 0.2 mL of the supernatant was transferred to HPLC vials for concentration determination by HPLC-ELSD (see supplementary materials). The concentration of PS80 left in the supernatants [*PS*80]_Ins_PS80_ was corrected for evaporation by the following equation:

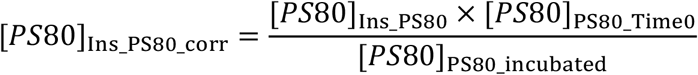

Finally, the molar PS80:insulin ratio of the spherulites was calculated as:

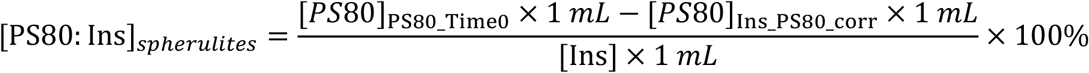

Where [Ins] represents the concentration of insulin (mM) in the spherulites dispersed in the liquid, and [PS80] is the concentration of PS80 (mM) remaining in the same spherulites solution.

### 2.6 Molecular dynamics (MD) and Pulling Simulations

Full details of the various MD simulations are given in the supplementary materials.

#### 2.6.1 Insulin-PS80 Simulation in 0.5 M NaCl Solution at pH=2

The structure of insulin was derived from X-ray diffraction (Whittingham et al. 2002) as deposited in the PDB database (PDB entry 1GUJ). The Amber force field was used (Wang et al. 2000). The PS80 was modelled using the GAFF2 (Wang et al. 2004, Wang et al. 2006) force field with Restrained ElectroStatic Potential (RESP) atomic charges (Bayly et al. 1993) found using the ACPYPE program (Sousa da Silva and Vranken 2012). The simple point-charge (SPC) (Berendsen et al. 1981) water model was used. A single isomer of PS80 was constructed. 20 monomers of insulin and 20 PS80 molecules were randomly positioned in a cubic simulation box of 20 × 20 × 20 Å^3^ and embedded in water, including 0.5 M NaCl. Periodic boundary conditions were invoked. After initial energy minimization and equilibration, 17 ns of MD simulation were conducted at constant temperature and pressure.

#### 2.6.2 PS80-induced Insulin Dimer Free Energy by Umbrella Simulation

Two different simulations were performed to study the stability of insulin dimers in contact with PS80. The force fields used for two simulations were the same as in section 2.6.1. The starting point for these simulations were obtained in the following way: a) Insulin dimer without PS80; the insulin dimer was obtained from the study of Whittingham et al. (Whittingham et al. 2002) and the protonation adjusted to pH 2 by using the pKa values obtained from the PropKa server (Olsson et al. 2011). b) Insulin dimer, similar to the one in simulation (a), but with one PS80 molecule attached to insulin on the opposite side of the insulin-insulin interface. The aggregated insulin-PS80 system was retrieved from the endpoint of the insulin-PS80 aggregation simulation described in 2.6.1.

In both cases, the system was allowed to equilibrate similar to the method used in 2.6.1, in a cubic box with periodic boundaries wherein there was SPC water and 0.5 M NaCl including neutralizing counterions. Following this equilibration, the system was simulated at 300 K for 10 ns, under which the assemblies remained stable.

The pulling-simulations and umbrella sampling were performed as follows. Structures from the end of each of the trajectories were used as starting points. The aggregates were placed in a rectangular box with dimensions sufficient to satisfy the minimum image convention and provide space for pulling dimension to take place along the z-axis. The insulin dimer was placed so that the pull along the z-axis was perpendicular to the plane of the dimer interface. As before, SPC water was used to represent solvent, and 0.5 M NaCl was present in the simulation cell. Equilibration was performed for 100 ps under an isothermal-isobaric ensemble at constant concentration (NPT ensemble), using the same methodology as described above.

Following equilibration, restraints were removed from all molecules except insulin monomer A; this was used as an immobile reference for the pulling simulations. For each of the simulations, insulin monomer B was pulled away from monomer A over 500 ps, using a spring constant of 1000 kJ mol^-1^ nm^-2^ and a pull rate of 0.01 nm ps^-1^. A final center-of-mass (COM) distance between monomer A and B of approximately 5.5 nm was achieved. From these trajectories, snapshots were taken to generate the starting configurations for the umbrella sampling windows.

The number of windows sampled for each simulation was 20, sampled asymmetrically to ensure a thorough sampling of the configurational space during the dissociation of the aggregate. In each window 10 ns of MD was performed. Analysis of the results was performed with the weighted histogram analysis method (WHAM) (Hub et al. 2010) as implemented in the GROMACS 5.1.1 software package (Abraham et al. 2015). Binding free energies were derived by analysis of the one-dimensional potential of mean force (PMF) curves along the pulling direction. A boot-strap method was used to assess the error of the derived free energies.

### 2.7 Optical Microscopy

#### 2.7.1 Polarized Light Microscopy

10 µL of the spherulites sample were placed on a glass slide without further dilutions. The polarized light microscopy images were obtained on a Leica DMi8 optical microscope using a 20× objective (Leica Microsystem, Wetzlar, Germany).

#### 2.7.2 Confocal Microscopy

Spherulites samples were diluted 1:20 (final insulin concentration of 0.15 mg/mL) and stained with ThT with a final concentration of 80 µM. 200 µL of stained samples were placed on microscope chamber slides and imaged at 1024 × 1024 pixels resolution using a Leica TCS SP5 confocal laser scanning microscope, using a 40×/1.25 oil objective (Leica Microsystems, Wetzlar, Germany). Confocal imaging was performed using λ_ex_=470 nm (Leica “white light” laser) and fluorescence was detected in the range λ_em_=485-550 nm.

### 2.8 Fluorescence Lifetime Imaging Microscopy

Sample preparation was identical to that described for samples for confocal imaging. Fluorescence lifetime imaging measurements were acquired in the time domain by means of a picoHarp 300 standalone TCSPC module (PicoQuant, Berlin, Germany). 256 × 256 images were acquired at a scanning frequency of 400 Hz and Leica “white light” laser was used to excite ThT (λ_ex_=470 nm, λ_em_=485-550 nm) (De Luca et al. 2020, Fennema Galparsoro et al. 2021)

Data analysis was performed by means of the SimFCS 4 program (Laboratory for Fluorescence Dynamics, University of California, Irvine, CA, available at www.lfd.uci.edu). FLIM calibration of the system was obtained by measuring the known lifetime of fluorescein, which shows a single exponential decay with characteristic lifetime of 4.0 ns (Wang et al. 2013).

A “phasor analysis” was performed for FLIM measurements, see details in (Digman et al. 2008, Ranjit et al. 2018). Briefly, it is a Fourier domain technique, which allows the transformation of the fluorescence decay from each pixel in the image to a point in the phasor plot. All possible single exponential lifetimes lie on a 0.5 radius semicircle going from point (0, 0) to point (1, 0), corresponding to τ=∞ and τ=0, respectively. As phasors follow the vector algebra, a multicomponent decay will lie inside the semicircle and is the result of the linear combination of the single exponential lifetime components. As an example, a double exponential decay would lie within the universal circle on the line connecting the two single exponential components.

### 2.9 Fourier Transform Infra Red Microscopy (Micro-FTIR)

Insulin spherulites were prepared as described in section 2.2. Spherulites in microtubes were isolated by centrifugation at 9,279 g, 10 °C for 10 min. The supernatant was carefully discarded, and the pellet washed with D_2_O to remove remaining H_2_O and salt. The samples were centrifuged again, the supernatant removed and replaced by 1 mL of fresh D_2_O, and finally kept in D_2_O overnight to allow H-D exchange.

Micro-FTIR measurements were performed at RT using a LUMOS Fourier transform infrared microscope (Bruker, Billerica, MA, USA), equipped with a photoconductive MCT detector with liquid nitrogen cooling. Visual image collection was performed via a fast digital CCD camera integrated in the instrument. Aliquots of aggregated samples in D_2_O were placed between two CaF_2_ windows separated by a 17 μm Teflon spacer, ensuring a common path length for all absorption measurements. FTIR spectra were acquired in transmission mode between 4000 and 700 cm^−1^, with a spectral resolution of 2 cm^-1^. Each spectrum represents an average of 128 scans in a 25 μm × 25 μm size region of interest. Spherulites with higher density were not measurable in the present configuration.

### 2.10 Scanning Electron Microscopy

The imaging was performed on a Quanta™ 3D FEG scanning electron microscope (Thermo Fisher Scientific, Hillsboro, OR, USA). After 42 hours of incubation, spherulites were pelleted after centrifugation for 10 min at 9,279 g and 10 °C. The supernatant was removed, and 1 mL of deionized water was added to resuspend the spherulites. A second cycle of centrifugation (10 min at 9,279 g and 10 °C) and subsequent removal of supernatant was used to wash the spherulites. The obtained highly concentrated suspension in the pellet was taken out with a plastic pipette and placed on a glass slide for drying overnight at ambient conditions. Dried samples were then mounted on carbon tapes and sputter coated with 0.2 nm gold with Leica EM ACE200 (Leica Microsystems, Wetzlar, Germany) before imaging. Images were acquired at an acceleration voltage of 2 kV. Void interspaces were observed on the surface of the spherulites. The areas of these void interspaces was quantified based on SEM images of the spherulites at magnification of 6500× and at least seven spherulites for each sample were analyzed. The area was determined by image J, according to the brightness difference between dark void interspaces and the surroundings. Areas were calculated by automatic counting of the pixels constituting the dark void interspaces. The number of pixels for the dark spots was then transformed into an area depending on the spatial resolution of the specific micrograph. Significant irregularities on the spherulites surface were avoided by manual zone selection. The void interspaces were separated into different groups (bins) with a bin width of 0.1 µm^2^ between 0.05 and 6 µm^2^ (first bin 0.05-0.1 µm^2^; last bin 5.9-6.0 µm^2^). Fractional distribution was calculated from the *A/A*_*s*_ ratio, where *A* is representing the total area of void interspaces of a specific bin, and *A*_*s*_ the total area of all void interspaces. An example is shown in supplementary materials Figure S2.

### 2.11 Spherulites Dissociation and Release of Soluble Insulin molecules

The procedure to determine release of soluble insulin from the spherulites is schematically shown in Figure S3. 1 mL of each spherulite sample (3 mg/mL insulin aggregated with different concentrations of PS80) was centrifuged twice for 10 min at 9,279 g and 10 °C. After the 1^st^ centrifugation step, the supernatant was discarded and 1 mL purified water was added to resuspend the spherulites. After the second cycle, the supernatant was removed and 1 mL 0.1 M boric acid-NaOH (pH 9) buffer was added. The buffer was prepared by adding 309 mg boric acid and 65 mg NaOH in 50 ml purified water, pH was adjusted by using 5N HCl and 5N NaOH. The pH value of the solutions was checked after washing. Spherulites in the pH 9 buffer were kept at RT and, at different time points within 120 hours, the sample was centrifugated (10 min at 9,279 g and 10 °C) and 100 µL supernatant out of the total volume of 1 mL was withdrawn for concentration determination of insulin released from the spherulites. 100 µL buffer was added after each time point of sample withdrawal to reestablish the original volume of the solution. Spherulites samples were resuspended by gently turning the microtubes up and down. Insulin concentration was determined by Nanodrop 2000C as previously mentioned. Experiments were repeated on two different batches of spherulites for each sample and three replicates for each time point were analyzed.

## 3. Results and Discussion

### 3.1 PS80 Affects the Kinetics of Insulin Spherulite Formation

As a first step, we monitored the insulin aggregation kinetics at different PS80 concentrations using ThT fluorescence. ThT is known to increase its quantum yield when in complex with aggregates of amyloid origin (Stsiapura et al. 2008).

As shown by the normalized ThT kinetics (Figure 1a, kinetics prior normalization can be found in Figure S4 in SI), the presence of PS80 significantly affected the aggregation kinetics by modifying both the lag time and the growth rate in a concentration-dependent manner. In absence of PS80, the lag time was approx. 2 hours, followed by a rapid increase of the ThT signal and a plateau phase. At 0.52 mM and 5.17 mM PS80, the lag time was > 5 hours, and the further growth of the signal proceeded over a period of several hours, showing that the presence of the surfactant also affected the growth rate. To obtain a model-free rate constant of the process, the inverse of the time at which the reaction reached 50% of its completion (1/t_50%_) was determined. In Figure 1b, both the 1/t_50%_ and the surface tension of the insulin-PS80 solutions are plotted as a function of the PS80 concentration. 1/t_50%_ (blue symbols in Figure 1b) drastically decreased at increasing PS80 concentration up to [PS80] = 0.05 mM with less pronounced, but still detectable changes for [PS80] > 0.05 mM. Interestingly, surface tension measurements (black symbols in Figure 1b) indicates that the CMC is approximately 0.05 mM. This may suggest a role of PS80 micelles in affecting the aggregation. Notably, we measured the residual insulin concentration not converted into aggregates after 42 hours of incubation (see methods) and estimated that >96% of the initial native insulin molecules was converted into aggregates at the tested PS80 concentrations (Figure 1c). Interestingly, in both the samples of highest and lowest concentration of PS80 tested, we verified by high performance liquid chromatography (see section 2.5) that the surfactant was incorporated into the aggregates. Specifically, we found that at the highest PS80 concentration, the molar ratio of PS80 to insulin in the spherulites was around 1:4.6 (5.1% (w/w)), and 1:30.3 (0.9% (w/w)) in the second highest PS80 concentration. It is worth to note that the ThT value at the plateau decreased at increasing concentration of PS80 (see Figure S4 in SI). Such an effect can be determined by a number of factors, including specificity and accessibility of ThT binding sites, and the macroscopic properties of the binding site environment.

**Figure 1.**
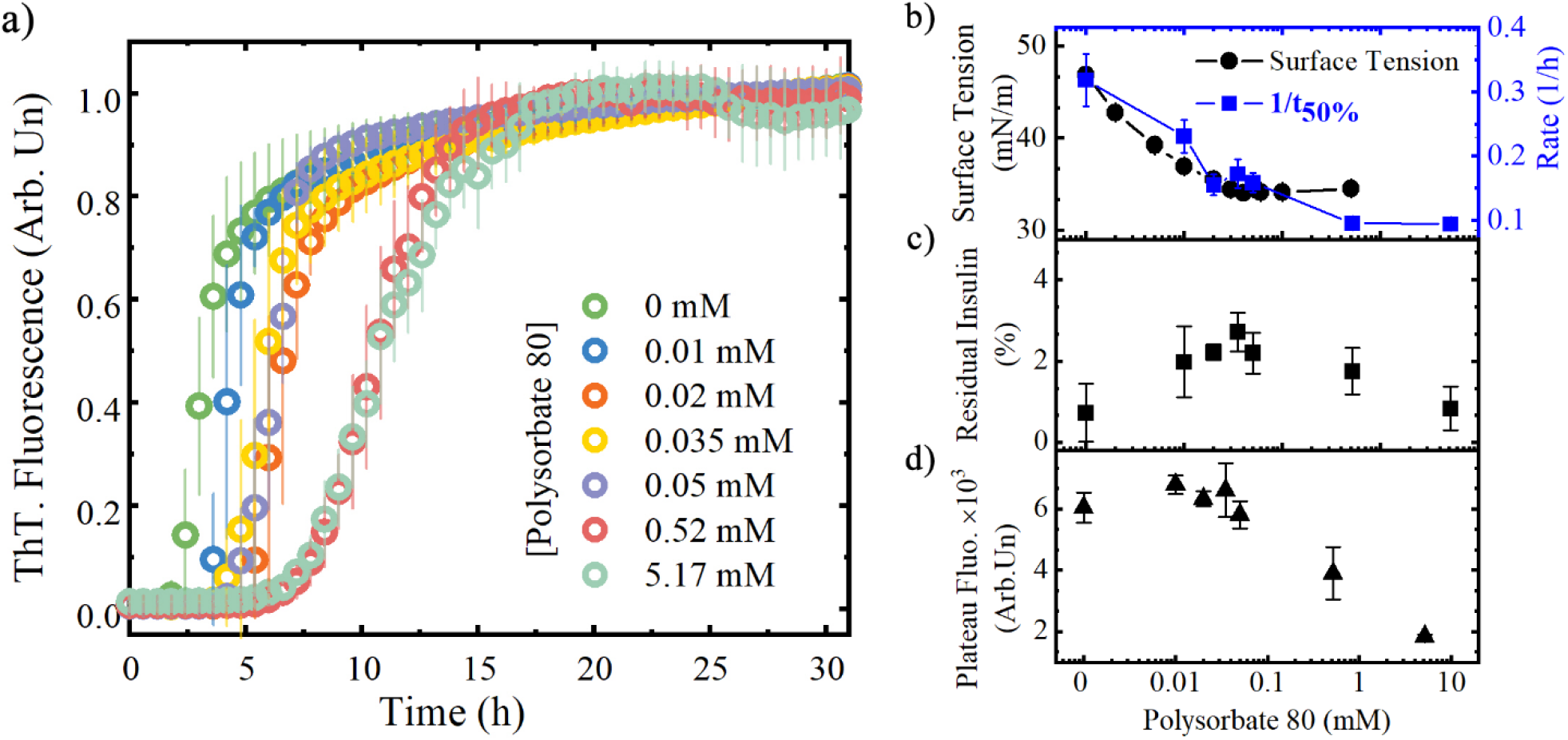
Kinetics of spherulite formation: (a) Thioflavin T (ThT) fluorescence kinetics for insulin at different concentrations of PS80 (Mean ± SD, n=4). 3 mg/mL insulin was dissolved in 20 %(v/v) acetic acid, 0.5 M NaCl solution, pH 1.7, at different PS80 concentrations and incubated at 45 °C for 42 hours. ThT concentration was 20 µM. Measurements are normalized to the plateau value. (b) Surface tension measurements of different concentrations of PS80 in 20% (v/v) acetic acid, 0.5 M NaCl, pH 1.7 in the absence of insulin (black circles, Mean ± SD, n=3) and the rate of aggregation 1/t_50%_ (blue squares, Mean ± SD, n=4). Three samples of each PS80 concentration were measured for the surface tension measurements. The 1/t_50%_ was calculated according to the data from Figure 1. (c) Residual concentration of soluble insulin after 42 hours of incubation (Mean ± SD, n=3). (d) ThT intensity value at the plateau.

Taken together, the data in Figure 1 shows that PS80 affected the insulin amyloid aggregation kinetics, causing a significant increase in the lag time, in particular at concentrations higher than the CMC. PS80 did not change the conversion efficiency of native protein into aggregates, but it was incorporated into the inner structure and/or onto the surface of the aggregates. Moreover, spherulites containing PS80 may provide a modified environment for Thioflavin T (e.g. increased the rigidity), resulting in different quantum yield of the dye and fluorescence values at the plateau of the kinetics (see Figure 1d and Figure S4 in supplementary materials). Notably, the addition of PS80 to the starting solutions did not change the secondary structure of insulin. There is no significant difference detected by CD, with all the spectra showing a native α-helix structure profile prior to incubation (Figure S5). Similarly, no differences were detected in the near-UV CD region (data not shown).

### 3.2 PS80 Affects Insulin-Insulin Interactions

The pronounced effect of PS80 on the lag time as displayed in Figure 1 is likely connected to the ability of insulin molecules to effectively interact with each other, either by direct PPIs or PSIs (Choi et al. 2014, Månsson et al. 2014, Arosio et al. 2015, Holubová et al. 2021). Indeed, a similar effect was previously reported for insulin in presence of ethanol, with an increase of the lag time due to the decrease of the dielectric constant of the medium and an ethanol-induced change of the protein hydration (Vetri et al. 2018).

We used molecular dynamics simulations to investigate the molecular interactions and binding between insulin molecules in presence and absence of PS80. Specifically, comparison of two configurations corresponding to insulin solution and insulin in presence of PS80, respectively, was done. Mixtures of 20 insulin molecules with 20 PS80 molecules correspond to the experimental condition of 0.52 mM PS80. For an insulin system in absence of PS80 and within a simulation time range of 20 ns, we only observed the occurrence of monomers (Figure 2a).

**Figure 2.**
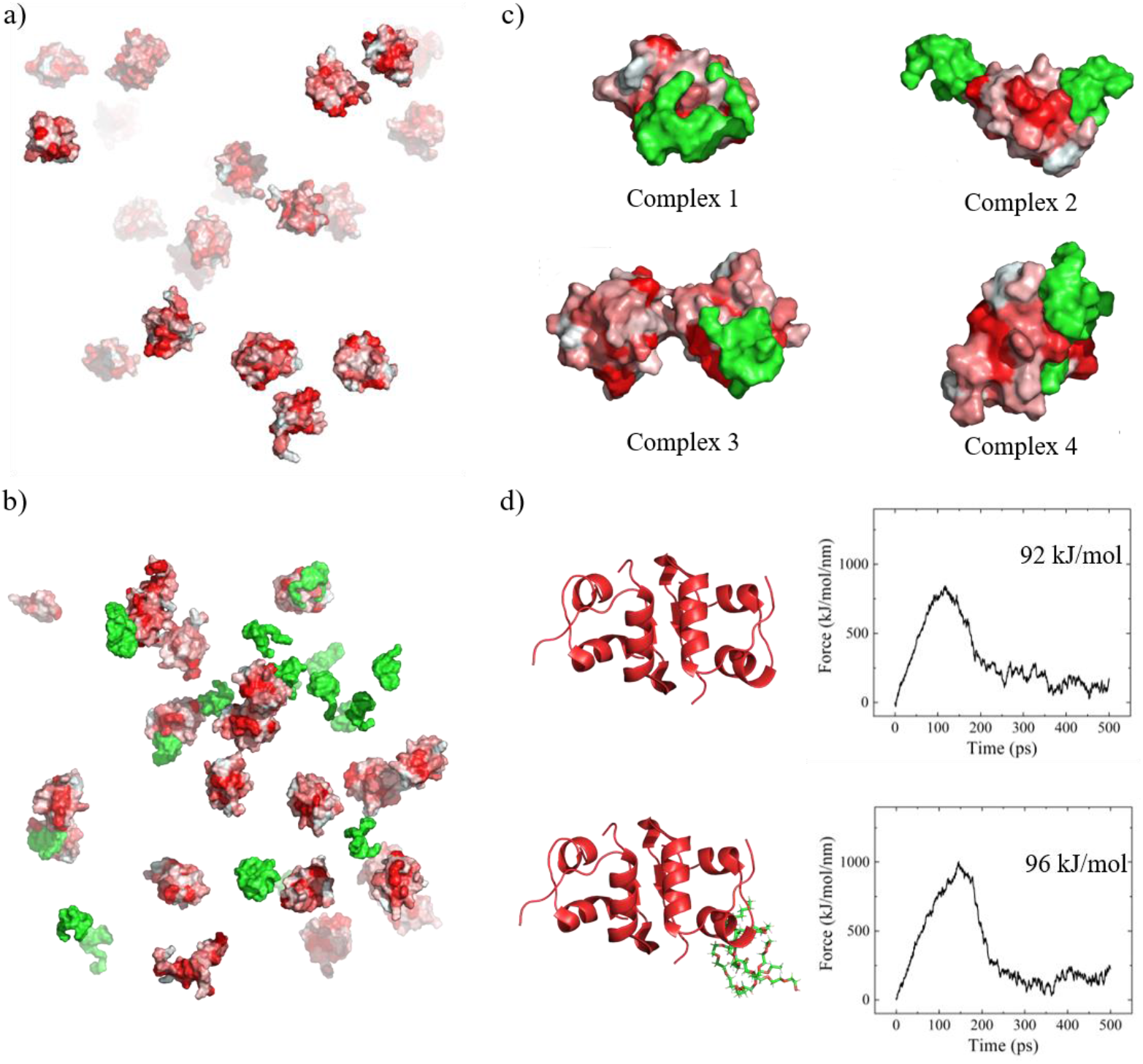
Simulations of (a) PS80-free insulin ensemble and (b) 20 insulin molecules in presence of 20 PS80 (green) molecules. (c) Complexes created from the simulation in b). Red color on the protein surface in (a), (b) and (c) indicates hydrophobic region of the molecules. (d) Pulling experiment by molecular docking. The pulling simulation on the dimers is performed by immobilizing one insulin monomer, while the other is pulled away using a harmonic force. The PS80 molecule is not constrained during the pulling experiments. The free energies of interaction were estimated based on a weighted histogram analysis method (WHAM) of sampled configurations along the reaction coordinate. The error associated with the energy minima is ±4 kJ/mol for each system.

Interestingly, a different scenario occurs when PS80 is added. PS80 interacts with insulin, generating four different complexes (Figure 2b and 2c) with PS80 mainly binding to insulin hydrophobic residues (see Table S1 in supplementary materials). Moreover, we note that insulin tends to form dimers in presence of PS80 within the timescale of the simulations (see complex 3 in Figure 2c). These dimers have a small interaction interface, and are therefore rather weakly bound compared to the dimers observed in the crystalline state (Whittingham et al. 2002). We hypothesize that the formation of strongly bound dimers, in agreement with the crystal dimers, will take place at longer timescales inaccessible by our atomistic simulations. For such strongly bound dimers, corresponding to the crystal dimer, we calculated the free energy needed to separate the two insulin molecules (Figure 2d), retrieving information on the binding energies between insulin molecules. For insulin dimers in absence of PS80, 92 kJ/mol (37 kT) is needed for the dissociation. This value is barely altered when PS80 is bound on the external part of one of the monomers within the dimer (96 kJ/mol or ∼39 kT).

The MD simulations suggest that the presence of PS80 generated a more heterogeneous sample with a propensity to form dimers. Insulin amyloid self-assembly at acidic pH values is considered to proceed via aggregation between monomers (Brange et al. 1997, Frankær et al. 2017), while formation of dimers is considered to increase the stability of insulin against fibril formation (Vinther et al. 2012). Thus, the increased tendency to form dimers with increasing PS80 concentration may explain the observed delay of the aggregation process (Figure 1a). It is also worth noting that insulin aggregation is known to be highly catalyzed in presence of different types of surfaces (Foderà et al. 2009, Li and Leblanc 2014). When adsorbed to an interface with high surface tension, proteins can increase their exposed area in contact with the surface (Randolph and Jones 2002), giving rise to nucleation points from which aggregation may develop (Nayak et al. 2008). The addition of surfactant decreases the surface tension, reducing protein adsorption (Randolph and Jones 2002) and in turn the catalytic effect of surface. Reduced adsorption may thus also have contributed to the delay of the aggregation process.

### 3.3 PS80 Affects the Morphology of Insulin Spherulites

To investigate the effect of PS80 on the morphology of the aggregates, we performed cross-polarized microscopy analysis of the aggregated samples (Figure 3a). The characteristic Maltese cross-pattern is detected for all the samples, confirming the presence of spherulites. Spherulites are also readily detectable via confocal microscopy using ThT staining (Figure 3b).

**Figure 3.**
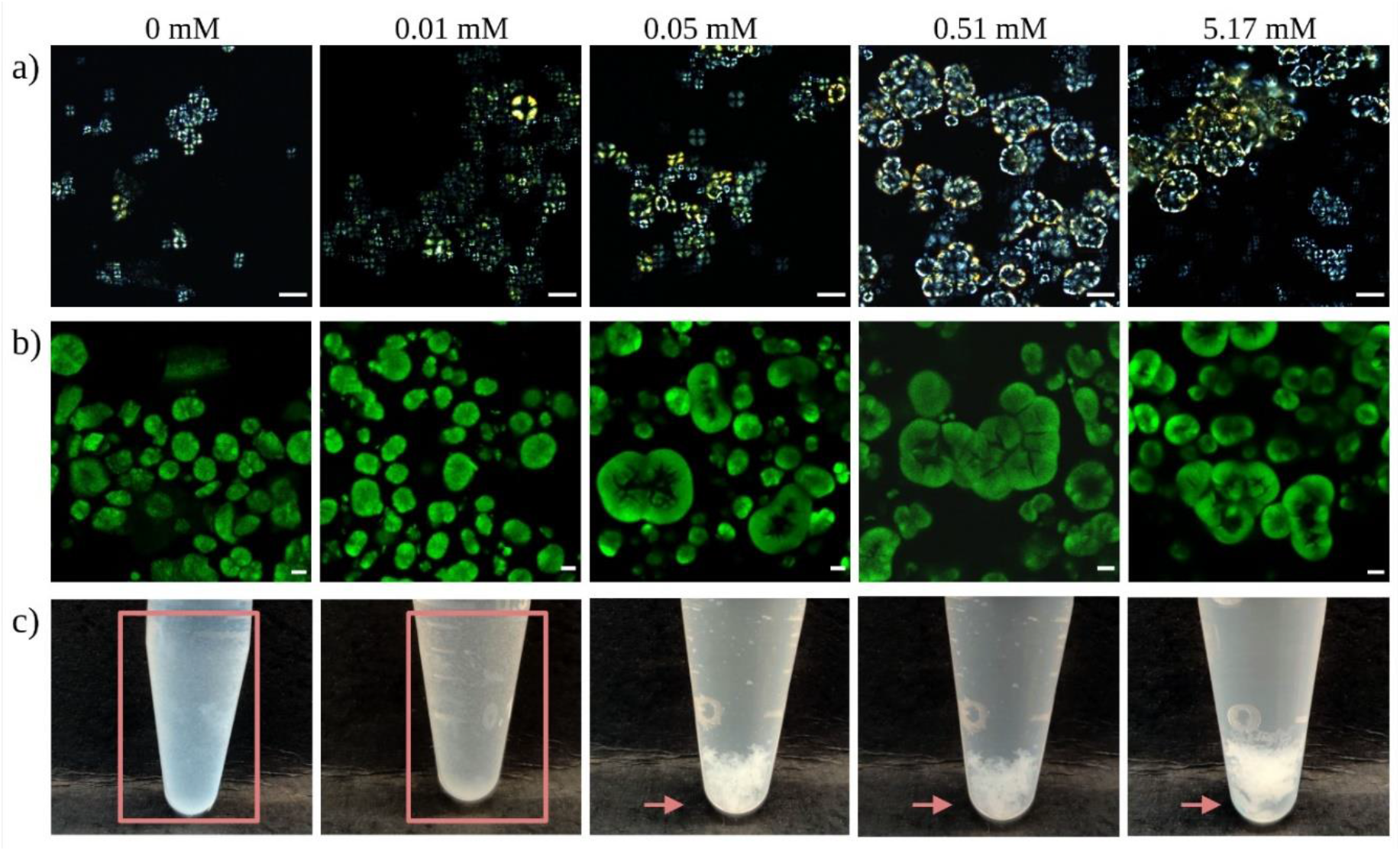
Morphology of the aggregates produced from solution of 3 mg/mL insulin in 20 % (v/v) acetic acid 0.5M NaCl pH 1.7, at different PS80 concentrations and incubated at 45 °C for 42 hours. Numbers on top of the figures indicate the concentration of PS80 added in the insulin solution. a) Polarized light microscopy (scale bar: 100 µm), and b) confocal microscopy of spherulites stained with Thioflavin T (scale bar: 20 µm). c) Spherulites in microtubes after 42 hours incubation. Arrows indicate the sedimentation phenomenon in presence of PS80 ≥ 0.05 mM.

Both in Figure 3a and 3b, in absence and at low concentrations of PS80, individual spherulites are observed within the samples. The scenario changes when [PS80] ≥ 0.05 mM (above CMC), where spherulites tended to form either clusters or large mesoscopic agglomerates that extended up to hundreds of µm (Figure. 3a and b). Furthermore, by visual inspection of the microtubes, spherulites formed at [PS80] ≤ 0.01 mM (below CMC) appeared to be distributed on the wall of the microtubes, while this phenomenon was less pronounced for [PS80]≥ 0.05 mM. At [PS80]> 0.05 mM (above CMC and in correspondence to the formation of spherulite clusters), the aggregates tended to sediment (Figure 3c). The presence of PS80 likely reduces the surface tension of the interface between solution and microtube walls (Khan et al. 2015). In our study, this phenomenon likely suppressed protein adsorption and the consequent formation of highly concentrated insulin areas in proximity of the surface. This would reduce the influence of surface-catalyzed phenomena in the aggregation process, and nucleation in the bulk becomes more significant than at lower PS80 concentrations. This apparently leads to the formation of larger µm-sized clusters in the bulk, which readily sedimented (arrows in Figure 3c).

To further investigate the morphological features of spherulites and the surface properties of the aggregates, we performed scanning electron microscopy (SEM). Results of SEM measurements are reported in Figure 4.

**Figure 4.**
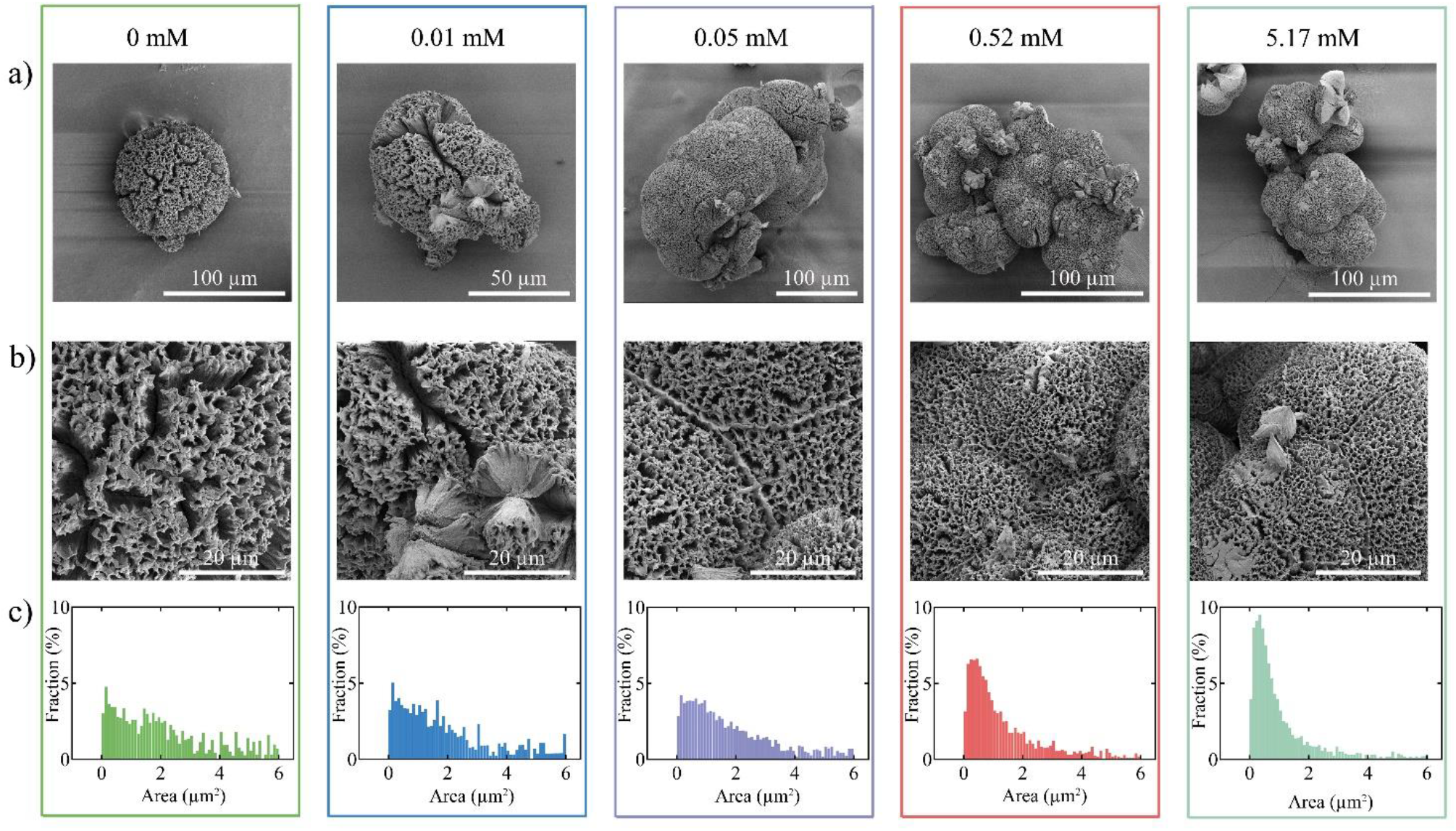
a-b) Scanning electron microscopy (SEM) images of spherulites produced from a solution of 3 mg/mL insulin in 20 %(v/v) acetic acid 0.5M NaCl, pH 1.7, at different PS80 concentrations and incubated at 45 °C for 42 hours. Numbers on top of the figures indicate the concentration of PS80 added in the insulin solution. b) Higher magnification of selected areas from a), acquired using the same magnification (6500×). c) Size distribution of void interspaces on spherulites surface. Seven areas containing spherulites for each condition were chosen for imaging, while different areas on each spherulite were used for size distribution calculation. >1000 void interspaces were included in the size distribution analysis. The area of interspaces was determined by evaluating the dark (black) areas on each spherulite.

Isolated spherulites with a pronounced roughness of the surface were observed in absence of PS80. In contrast, in presence of PS80 at concentrations higher than 0.05 mM, a pronounced clustering among the spherulites was observed and the spherulite surface appears smoother (Figure 4a). This was confirmed by a quantitative analysis of surface roughness. We evaluated the roughness as the distribution of void interspaces on the surface (Figure 4c). A broad flat distribution, spanning from few tens of nm^2^ to six µm^2^, characterized the surface of the spherulites formed in absence of PS80 (Figure 4c). In contrast, at increasing PS80 concentrations spherulites presented a surface with void interspaces that peak around 500 nm^2^ (Figure 4c), confirming the marked difference that can qualitatively be appreciated from the SEM images (Figure 4b). This may indicate a higher packing density of proteins within the spherulites formed in the presence of PS80.

### 3.4 PS80 Affects the Molecular Structure of Insulin Spherulites

Data in Figure 4 support the idea of a PS80-induced modification of the morphology of the spherulites, which may reflect different molecular organization within the whole structure. To gain information on this aspect, we combined FLIM and the phasor approach using ThT and micro-FTIR. In Figure 5a the phasor plot obtained from measurements on samples at 0 mM, 0.01 mM, 0.035 mM, 0.05 mM, 0.52 mM and 5.17 mM PS80 concentration is reported. Phasor plot allows a graphic visualization of fluorescence decay at pixel resolution. Each position in the plot represents a characteristic lifetime and clouds of points in the phasor plot are generally identified as fluorescence lifetime distribution. Examples of qualitative analysis of individual samples are reported in the supplementary materials (see Figure S6)

**Figure 5.**
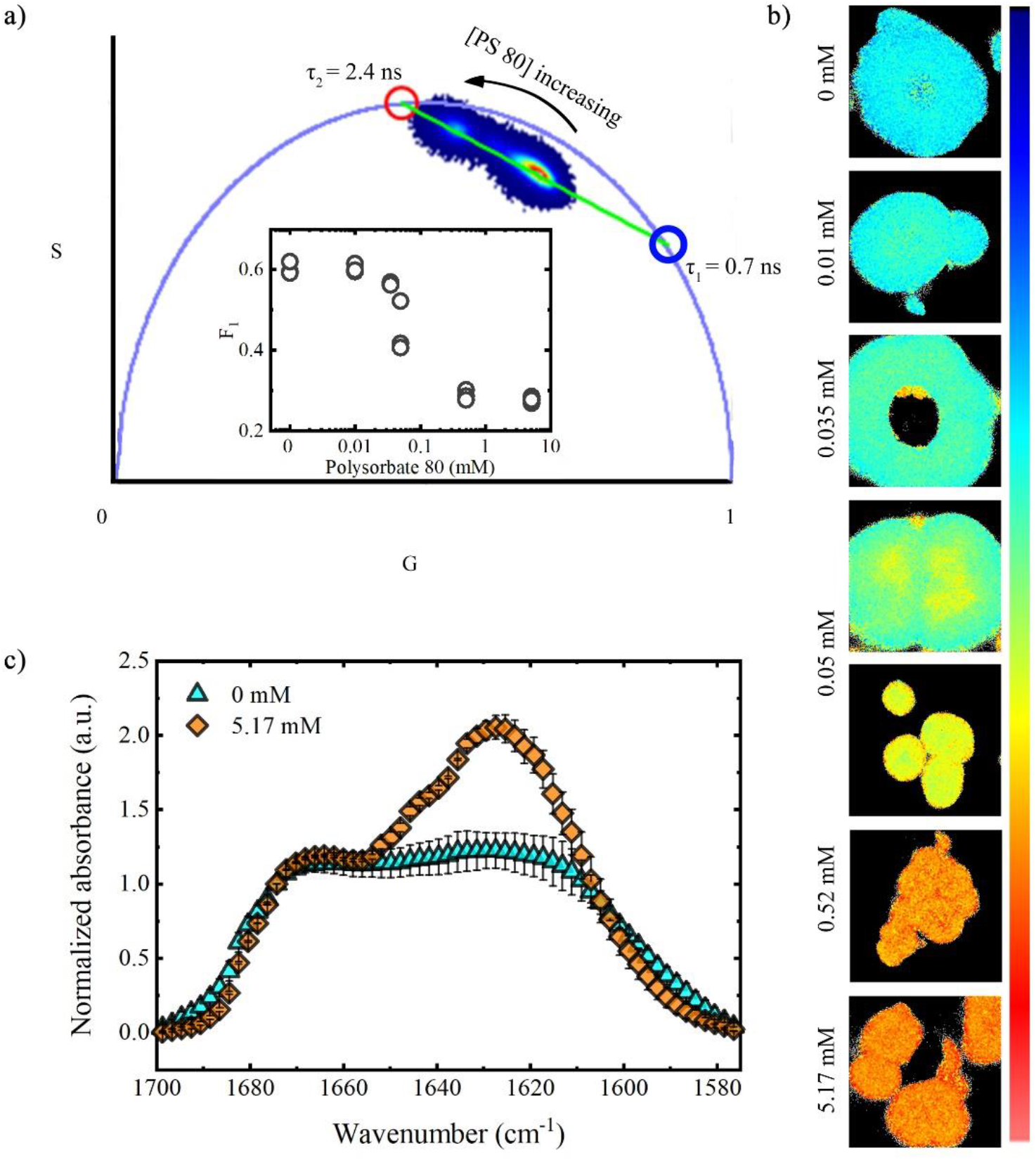
(a) Phasor plot extracted from FLIM measurements on insulin spherulites formed at different PS80 concentrations. Lifetime distributions clearly lie on a straight line (green) and for this reason are analyzed in terms of combination of two exponential decay. Characteristic lifetime components (τ_1_= 0.7 ns in blue and τ_2_= 2.4 ns in red) are identified from the intersection of the green line with the universal circle. Insert: F_1_ (fraction of τ_1_) as a function of PS80 concentration. (b) Lifetime fraction maps for the FLIM images at different concentration of PS80. Two images at CMC of PS80 are presented to highlight the heterogeneity of the sample. (c) FTIR spectra in the amide I band (1575 cm^-1^-1705 cm^-1^) of the sample without PS80 (cyan) and with 5.17 mM PS80 (orange). Data are normalized to the signal at 1675 cm^-1^. Three spherulites for each sample were measured (Mean ± SD, N=3)

In Figure 5a large lifetime distribution of ThT can be identified in the phasor plot, suggesting that the dye experiences three main distinguishable (see Figure S6 for better visualisation) and quite heterogeneous environments. Interestingly, the lifetime of ThT gradually increased with the addition of PS80. Measured lifetime distributions in the phasor plot clearly lie on a straight line connecting two points. This indicates the possibility of describing measured ThT fluorescence decays using a double exponential model.

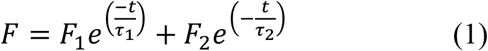

where the mono-exponential components (with characteristic lifetimes τ_1_ and τ_2_) can be identified via the intersection of the straight line with the universal circle. In our case we can identify two characteristic ThT lifetimes at τ_1_=0.7 ns (blue circle) and τ_2_=2.4 ns (red circle). Distances between each lifetime distribution and the single exponential phasors on the universal circle (F_1_ and F_2_) represent the fraction of each component (Digman et al. 2008, De Luca et al. 2020). In the insert in Figure 5a, we report the obtained F_1_ fractions as a function of the PS80 concentration and in Figure 5b representative 256 × 256 lifetime maps obtained for samples at different PS80 concentrations were color coded on a scale from blue (pure fast component, τ_1_ =0.7 ns) to red (pure slow component, τ_2_=2.4 ns). F_1_ dependence on the PS80 concentration clearly showed a biphasic behaviour. A high F_1_ value was found at PS80 concentrations below the CMC (∼0.05 mM) whilst at increasing surfactant concentrations the F_1_ value was reduced by more than 50% and remained constant at the highest concentrations of PS80. Importantly, Figure 5b reveals that spherulites formed at concentrations of PS80 above or below the CMC were characterised by a homogenous lifetime within the structure (see also Figure S6) and that only at/close to the CMC, different distributions of ThT fluorescence decays (two different lifetime distributions) coexists in the same sample (see also Figure S6).

Interestingly, double exponential decays were previously observed for ThT in different systems and related to peculiar properties of ThT environment with different rigidity (Stsiapura et al. 2008, Thompson et al. 2015) or to specific properties of the binding sites (Biancalana and Koide 2010, Sulatskaya et al. 2010, Lindberg et al. 2015, Ivancic et al. 2016, Sidhu et al. 2018). In a previous study using phasor approach for insulin spherulites samples, we found ThT fluorescence decays with same characteristic lifetimes as shown in the present study (De Luca et al. 2020). We attribute ThT lifetime changes within different aggregates structures to the superimposition of two main effects: the rigidity of the environment (due to the molecular-rotor nature of ThT (Stsiapura et al. 2008, Thompson et al. 2015) and the specificity of ThT binding site, which depends on the details of the β-sheet architecture. More specifically, we established a clear correlation between differences in ThT lifetime and peculiarities of intermolecular β-structures: higher ThT lifetimes correspond to higher content of β-structures (De Luca et al. 2020). Based on that, data in Figure 5a and 5b suggest that PS80 induced the formation of spherulites with higher content of intramolecular β-structures. In addition to this, different ThT binding modes and affinities may occur in spherulites formed at different PS80 concentrations, also affecting the observed ThT lifetimes. Stronger H-bonds and/or longer β-chains may indeed add more constraints and a higher rigidity to the ThT binding site with the outcome of inhibiting more effectively the non-radiative ThT rotation, eventually increasing the fluorophore’s lifetime.

The FTIR spectra acquired on single spherulites formed in the absence of PS80 (cyan) and at 0.51 mM PS80 (orange in Figure 5c) in the amide I band region (1575 cm^-1^-1705 cm^-1^) confirmed the difference in the secondary structure content. For both samples, the spectra were normalized to 1675 cm^-1^ and characterised by two main broad components centred at 1620 cm^-1^ and 1670 cm^-1^. The 1620 cm^-1^ peak is assigned to the aggregated β-sheets characteristic of the amyloid structure, while the peak at 1670 cm^-1^ represents the presence of α-helices, turns and loops (1660 cm^-1^) and antiparallel aggregated β-sheets (1670-1680 cm^-1^) (De Luca et al. 2020). Insulin spherulites formed at high concentration of PS80 had a higher content of aggregated β-sheets compared to the aggregates formed in absence of PS80, confirming the change in internal structure inferred from the FLIM analysis.

### 3.5 PS80 Affects the Stability of Insulin Spherulites in Alkaline Conditions

The previous sections show that the addition of PS80 changed the aggregation kinetics, morphology, and molecular structure of insulin spherulites, which may relate to differences in aggregate stability. Electrostatics forces are known to affect the stability of amyloid aggregates, and alkaline conditions promote the disassembly of insulin amyloid fibrils, leading to a release of soluble insulin molecules (Shammas et al. 2011, Santangelo et al. 2016). The stability of spherulites were therefore tested against critical pH change by dispersing them in an aqueous solution at pH 9 to achieve disassembly kinetics within a reasonable time frame.

Figure 6a displays the dissociation kinetics of insulin spherulites formed at different PS80 concentrations. Specifically, we monitored the concentration of soluble insulin released from the aggregates over time in boric acid buffer at pH 9 relative to the total insulin content in the spherulites. Independent of the PS80 concentration, spherulites show an initial fast release followed by a much slower sustained release of soluble insulin. Notably, the spherulites formed at [PS80]> 0.05M released soluble insulin species corresponding to ∼30-40% of the initial mass of spherulites, while a release up to ∼60% was detected for [PS80]<0.05M.

**Figure 6.**
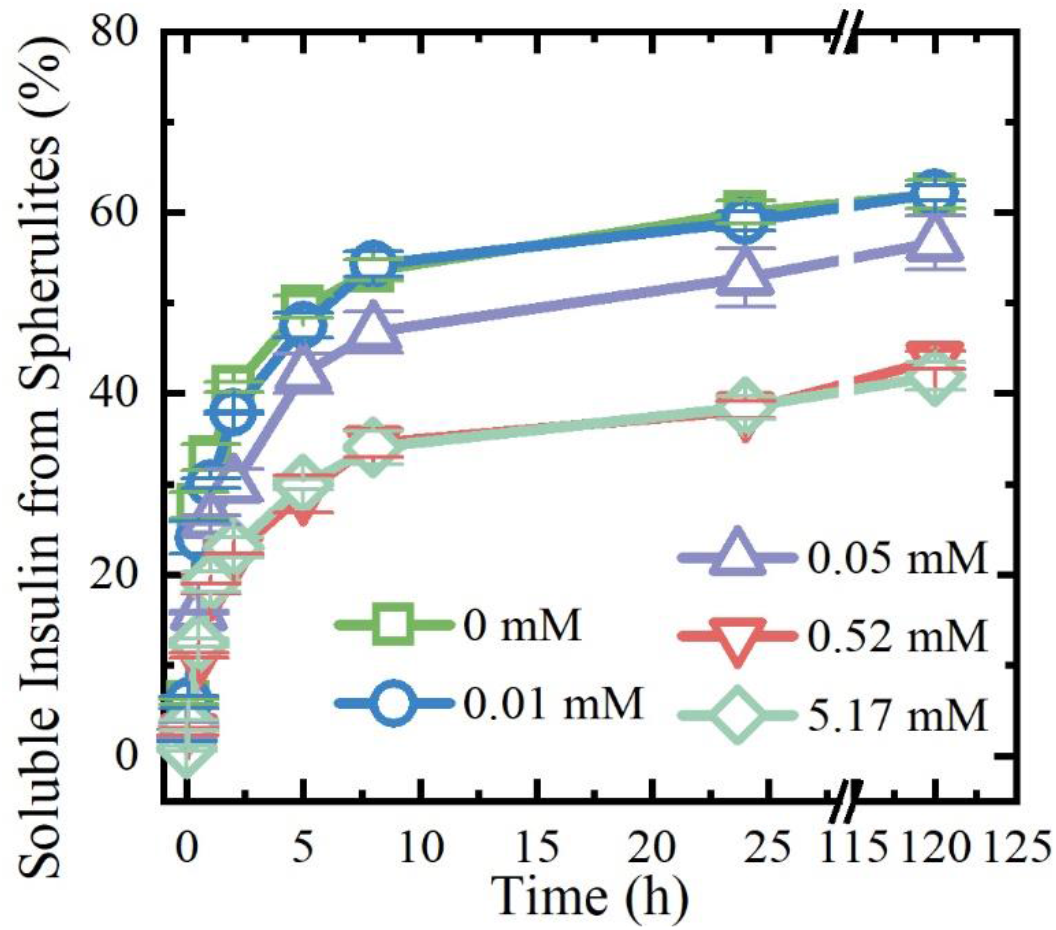
Stability of insulin-spherulites in pH 9 buffer. Dissociation and release of soluble insulin profiles for different insulin spherulites. The data is presented as the percentage of the amount of insulin released to the initial mass of insulin spherulites. Three spherulites samples of each PS80 concentration were included in the test (Mean ± SD, n=3).

Interestingly, a sustained release of soluble insulin species was observed up to 5 days (Figure 6a). These data suggest an enhanced stability of the spherulites formed at high PS80 concentration. No insulin release was observed within the same time-frame when the spherulites were suspended in buffers at pH values of 1.7, 5 and 7 (Figure S7).

When spherulites formed at pH 2 were exposed to pH 9, deprotonation of glutamic acid (pKa of side chain is 4.3), the carboxylic acid termini (pKa is 3.3), histidine (pKa of side chain is 6.5) and the N termini (pKa is 8.0) likely took place due to increased net negative charge (Pace et al. 2009, Shammas et al. 2011). This in turn will enhance the electrostatic repulsive forces between amino acids within the aggregates and likely contribute to the dissociation of spherulites. The deprotonation of the two N-termini on insulin appears to be crucial to facilitate disassembly and subsequent release, as minor to no release was observed even at pH 7. The enhanced stability for the spherulites containing high amounts of PS80 can be ascribed to the high degree of compactness of the structure (Figure 4) and the significant amount of hydrogen bonds involved in the β-aggregate structures.

## Conclusions

In this study, we present the effect of PS80 on the self-assembly mechanism, structure, morphology and stability of insulin amyloid-like spherulites. PS80 induced a delay in the onset of amyloid spherulites formation. This delay may be ascribed to the ability of PS80 to induce insulin dimers with increased stability against aggregation and to modify the surface properties of the sample holder, reducing surface-catalysed aggregation. Interestingly, the addition of this surfactant did not affect the total amount of insulin converted into aggregates. However, at increasing PS80 concentration, the level of clustering of amyloid spherulites was greatly affected, with a threshold around the CMC value for the surfactant. Moreover, when formed in solution with [PS80] > CMC, the spherulites acquired a very dense packing of the structure with an increase in the amount of β-sheet structure. This conferred an increased stability to the spherulites when exposed to alkaline conditions. Avoiding the appearance of protein particles in pharmaceutical products is one of the most challenging tasks in pharmaceutical sciences. The presence of surfactants like polysorbates is known to delay protein aggregation. Our findings point out that the addition of PS80 may alter not only the protein aggregation kinetics, but also the physicochemical properties of the protein aggregates. Such a change in properties is not obvious from single morphological assessments and requires a set of specific analytical methods. Further investigation is needed to evaluate whether the change in aggregate properties highlighted in our work has any impact on safety and efficacy. Importantly, using controlled self-assembly of biomolecular complexes has been widely discussed (Gupta et al. 2010, Tufail et al. 2018) including the possibility to exploit spherulites as carriers for drug delivery (Jiang et al. 2011). In this regard, our data may pave the way towards the use of co-solutes to alter the stability and morphology of the final structure, giving the opportunity to meet the different demands for target applications.

## Abbreviations

PS80: Polysorbate 80
CMC: Critical Micelle Concentration
FLIM: Fluorescence Lifetime Microscopy
SEM: Scanning Electron Microscopy
HPLC-ELSD: High Performance Liquid Chromatography with Evaporative Light Scattering Detector
FTIR: Fourier Transform Infra Red Microscopy

## Author Contributions

**Xin Zhou**: Investigation, Visualization, Formal Analysis and Writing - Original Draft. **Dirk Fennema Galparsoro**: Investigation and Writing - Original Draft. **Anders Østergaard Madsen**: Investigation and Writing - Review & Editing. **Valeria Vetri**: Project administration and Writing - Review & Editing. **Marco van de Weert**: Project administration and Writing - Review & Editing. **Hanne Mørck Nielsen**: Project administration and Writing - Review & Editing. **Vito Foderà**: Conceptualization, Funding acquisition, Writing - Original Draft.

## Acknowledgements

V.F. and X.Z. acknowledge China Scholarship Council (201709110108) for funding the project. V.F., D.F.G., and X.Z. acknowledge VILLUM FONDEN for supporting the project via the Villum Young Investigator grant “Protein Superstructures as Smart Biomaterials (ProSmart)” 2018−2023 (19175). The authors acknowledge the Core Facility for Integrated Microscopy, Faculty of Health and Medical Sciences, University of Copenhagen.). The authors thank Dr. Jijo Vallooran Joy (University of Copenhagen) for useful discussions about the experimental data. The authors thank Drug Research Academy that funded the NanoDrop 2000C, the Danish Research Council for Technology and Production Sciences for funding Centrifuge 5417R, and University of Copenhagen, together with Innovative Medicines Initiative Joint Undertaking (115363) that supported HPLC-ELSD system. The authors also kindly thank the VILLUM FONDEN (19175) for funding the CLARIOstar plate reader. and VILLUM FONDEN (19175), the Novo Nordisk Foundation (NNF16OC0021948) and Lundbeck Foundation (R155-2013-14113) for funding the Leica DMi8 microscope.

## Supplementary Materials

### High-Performance Liquid Chromatography (HPLC)-Evaporative Light Scattering Detection (ELSD)

The high-performance liquid chromatography (HPLC) system consisted of an Agilent 1200 Series HPLC system (Agilent Technologies, Palo Alto, CA, USA) equipped with a quaternary pump and auto-injector coupled to a 1260 Infinity evaporative light scattering detector (ELSD) (Agilent Technologies, Palo Alto, CA, USA). In-house nitrogen supply from University of Copenhagen was used as the source of the nitrogen gas for the ELSD. An HPLC column Gemini C18, 150 × 4.6 mm, 3 mm (Phenomenex, Torrance, CA, USA) was used in the experiment. The mobile phase A consisted of 5% (v/v) acetonitrile (VWR, ≥ 99.9 %, Radnor, PA, USA), while mobile phase B consisted of 95% (v/v) acetonitrile. Both phases also contained 0.1 %(v/v) trifluoroacetic acid (Merck, ≥99.0%, Saint Louis, MO, USA). The gradient conditions were as follows:

During the first 5 min of the analysis time, mobile phase B was maintained at 0%, and subsequently increased linearly to reach 60% at 6 min and 80% at 10 min. After maintaining 80% B from 10 to 15 min, the mobile phase was changed back to 0% B at 15.1 min, and the column was equilibrated until 20 min. Both the evaporator and the nebulizer temperature of the ELSD were maintained at 45 °C. The gas flow rate was set to 1.6 SLM.

The concentration of PS80 standard solution varied from 0.1 mg/mL to 1 mg/mL. The plot of log ELSD area versus log PS80 concentration showed a linear relationship (R^2^ > 0.99), the limit of quantification was 0.0067 mg/mL (0.051 mM).

### Molecular Dynamic Modulation Conditions

The insulin structure was obtained from the study of Whittingham et al. (Whittingham et al. 2002) as deposited in the PDB database (reference code 1GUJ). One monomer was used. The protonation of the insulin molecule was adjusted to pH 2 by use of the pKa values obtained from the PropKa server (Olsson et al. 2011), and the Amber force field was used (Wang et al. 2000). A single isomer of PS80 was used with x = 9, z = 3, y = 3, w = 5 according to the labeling system in Figure S1. The PS80 was modelled using the GAFF2 (Wang et al. 2004, Wang et al. 2006) force field with RESP atomic charges found using the ACPYPE program (Sousa da Silva and Vranken 2012). The SPC (Berendsen et al. 1981) water model was used. 20 monomers of insulin and 20 PS80 molecules were randomly positioned in a cubic simulation box of 20 × 20 × 20 Å, to which 0.5 M NaCl was added, including neutralizing counterions. Periodic boundary conditions were invoked.

The system was energy minimized and subsequently equilibrated in two steps, with position restraints applied to the peptide heavy atoms throughout. The first phase involved simulating for 100 ps under a constant volume (NVT) ensemble. Protein and nonprotein atoms were coupled to separate temperature coupling baths, and temperature was maintained at 310 K using the Berendsen weak coupling method (Berendsen et al. 1984). Following NVT equilibration, 100 ps of constant pressure (NPT) equilibration were performed, also using weak coupling (Berendsen et al. 1984) to maintain pressure isotropically at 1.0 bar.

Production MD simulations were conducted for 20 ns in the absence of any restraints. These MD simulations were performed using the GROMACS 5.1.1 software package (Abraham et al. 2015). The electrostatic and van der Waal forces were calculated with the particle mesh Ewald approach (Darden et al. 1993). The leap-frog algorithm was used as integrator together with the velocity rescaling thermostat (Bussi et al. 2007) and the Parrinello-Rahman barostat (Parrinello and Rahman 1981).

**Figure S1.**
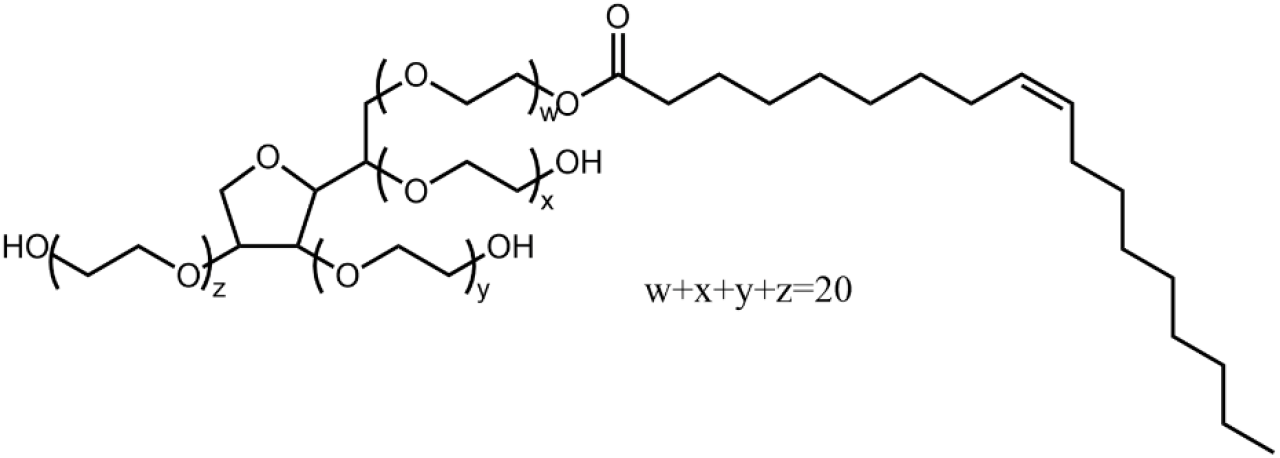
The chemical structure of Polysorbate 80

The 4 insulin and PS80 complexes that we obtained from the simulations are shown in Figure 2c of the main article. The residues of insulin interacting with PS80 are shown in Table S1.

**Table S1.**
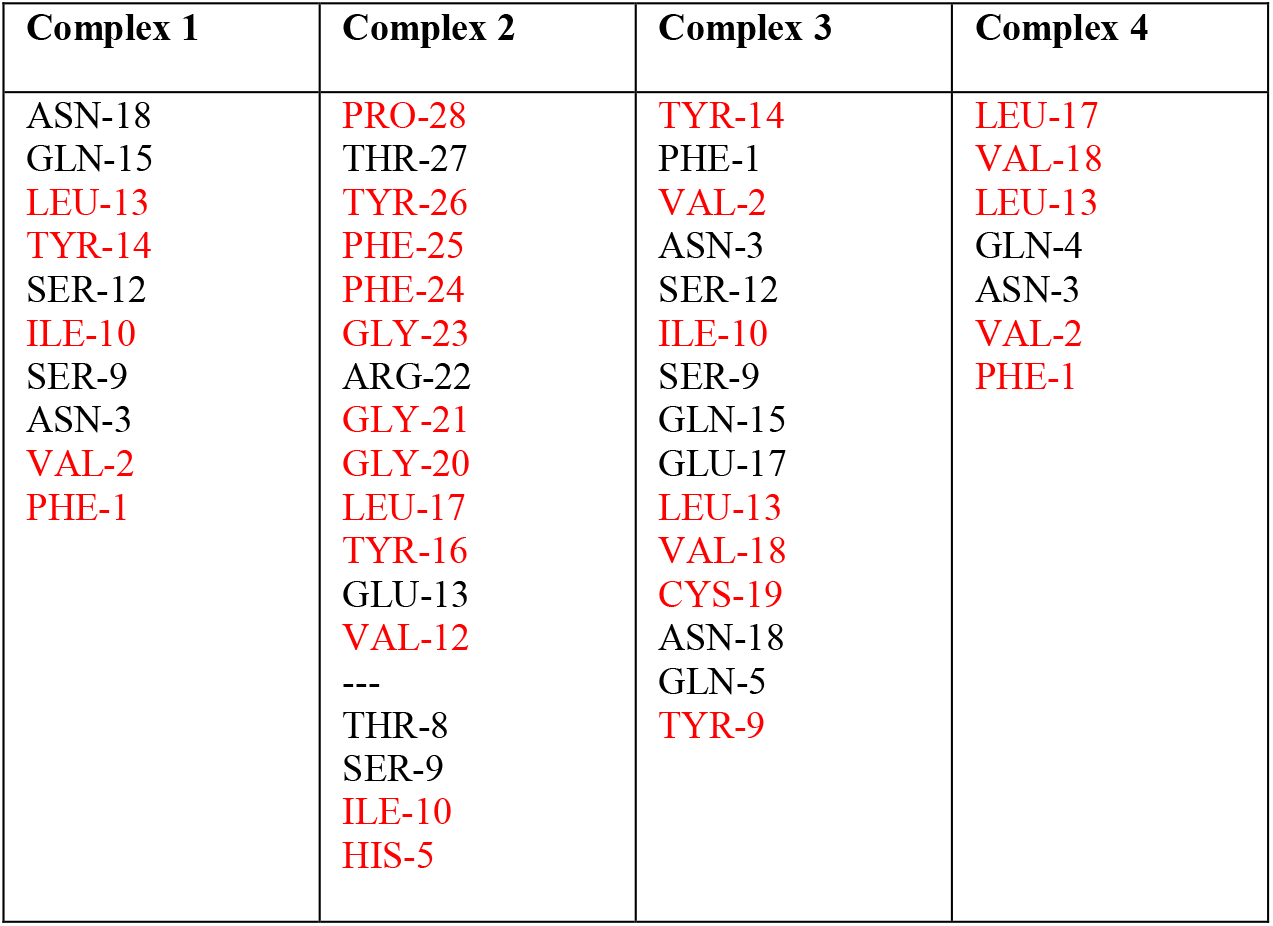
Residues of insulin interacting with PS80 in Figure 2c. Residues that are hydrophobic according to the hydrophobicity scale of Eisenberg et al (1984) have been colored red.

| Complex 1 | Complex 2 | Complex 3 | Complex 4 |
| --- | --- | --- | --- |
| ASN-18 | PRO-28 | TYR-14 | LEU-17 |
| GLN-15 | THR-27 | PHE-1 | VAL-18 |
| LEU-13 | TYR-26 | VAL-2 | LEU-13 |
| TYR-14 | PHE-25 | ASN-3 | GLN-4 |
| SER-12 | PHE-24 | SER-12 | ASN-3 |
| ILE-10 | GLY-23 | ILE-10 | VAL-2 |
| SER-9 | ARG-22 | SER-9 | PHE-1 |
| ASN-3 | GLY-21 | GLN-15 |  |
| VAL-2 | GLY-20 | GLU-17 |  |
| PHE-1 | LEU-17 | LEU-13 |  |
|  | TYR-16 | VAL-18 |  |
|  | GLU-13 | CYS-19 |  |
|  | VAL-12 | ASN-18 |  |
|  | --- | GLN-5 |  |
|  | THR-8 | TYR-9 |  |
|  | SER-9 |  |  |
|  | ILE-10 |  |  |
|  | HIS-5 |  |  |

### Determination of the Void Interspaces by ImageJ

ImageJ was used in this study to obtain the area of void interspaces on spherulites surface. The method was adjusted according to an example offered by ImageJ (for full detail, https://imagej.nih.gov/ij/docs/pdfs/examples.pdf), process details listed below:

1. Open an SEM image of spherulite with ImageJ (Figure S2a). Set the scale according to the scale bar on the image.
2. Change the image type to 8-bit. Adjust the threshold of the image. Use the slider bars to set the threshold range, until the red area only contains open spaces (Figure S2b).
3. Analyze – Analyze Particles. Set the size (µm^2^) as 0.05-6.00. Select “show outlines”. After clicking on “OK”, a list of areas of the chosen spaces should show, together with a drawing of all the chosen spaces (Figure S2c).

By following the steps above, void interspaces were selected. Analysis was performed on seven spherulites for each PS80 concentration. The number of spaces analyzed for each PS80 concentration is in the range 2000-14000.

**Figure S2.**
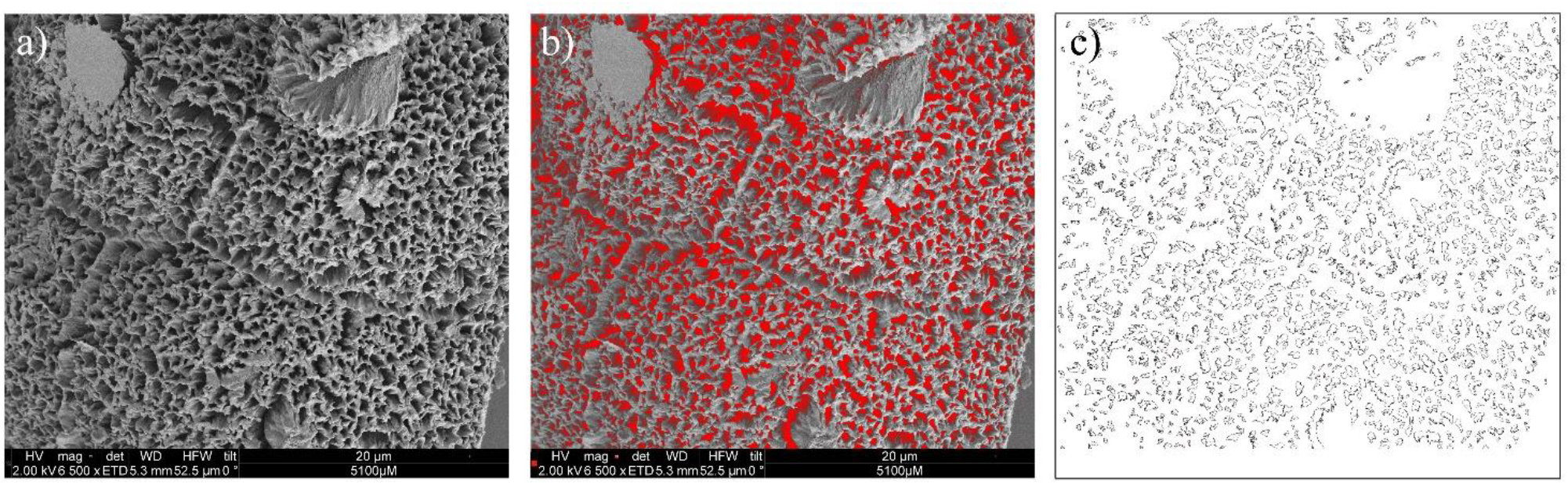
An example of using ImageJ to obtain the area of open spaces. (a) The original scanning electron microscopy (SEM) image of a spherulite. (b) Threshold adjustment of (a) for selecting the open spaces (in red). Outline of selected red area in (b).

### Protocol of Dissociation Study

The protocol of dissociation study in different pH values solutions were conducted as described below.

Spherulites washing steps:

- 1 mL insulin spherulites samples incubated with different amount of PS80 were centrifuged under the condition mentioned in Figure S3.
- The supernatant was carefully taken out and 1 mL purified water was added. The samples were resuspended and centrifuged again.
- The above two steps were repeated.

Steps for the measurement of soluble insulin dissociated from spherulites:

- After removing the water from the supernatant, 1 mL buffer (pH values were 1.7, 5, 7 and 9) were added.
- Spherulites in the buffer were kept at room temperature (RT) for 120 hrs. At 0, 1, 2, 4, 6, 8, 24, and 120 hrs, the spherulites samples were centrifuged under the specified conditions. 0.1 mL of the supernatant was taken out for measuring the insulin concentration. Each measurement was repeated 3 times, with 3 spherulites samples measured for each PS80 concentration. The average value was reported as the final result.
- During sampling 0.1 mL of fresh boric acid-NaOH buffer was added to the spherulites samples to reestablish the original volume. Samples were resuspended every time after the fresh buffer was added.

**Figure S3.**
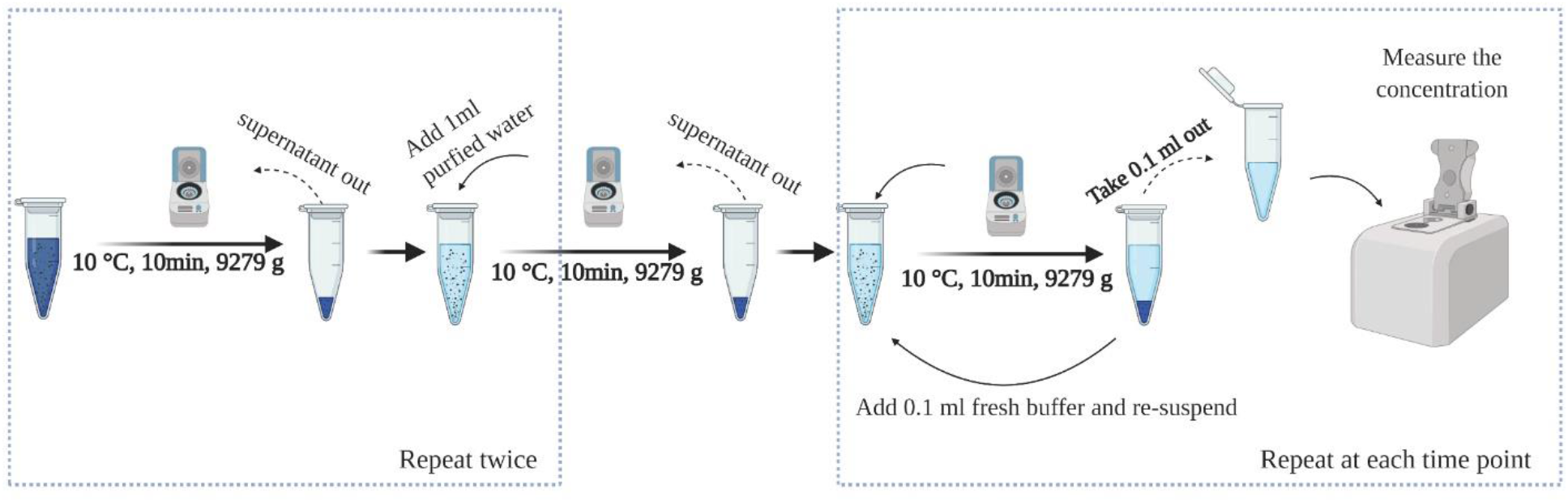
Schematic overview of the protocol for the dissociation of spherulites and release of soluble insulin study. Created with BioRender.com

### Thioflavin T (ThT) Kinetics

The ThT fibrillation kinetics without normalization is shown in Figure S4. As mentioned in section 2.3 of the main article, 3 mg/mL insulin was dissolved in 20 %(v/v) acetic acid and 0.5 M NaCl, pH 1.7, with increasing concentration of PS80. The insulin solutions were incubated in 96 well plates at 45 °C for 42 hrs. ThT was added using a final concentration of 20 µM. The addition of PS80 led to a longer lag time. Interestingly, a decrease of the maximum fluorescence was also observed (Figure S4). Upon PS80 addition, the plateau of the kinetic curves was reduced, especially when the concentration of PS80 was above 0.52 mM. This likely was due to fewer binding sites for ThT, as the total conversion of insulin into aggregates was similar for all samples.

**Figure S4.**
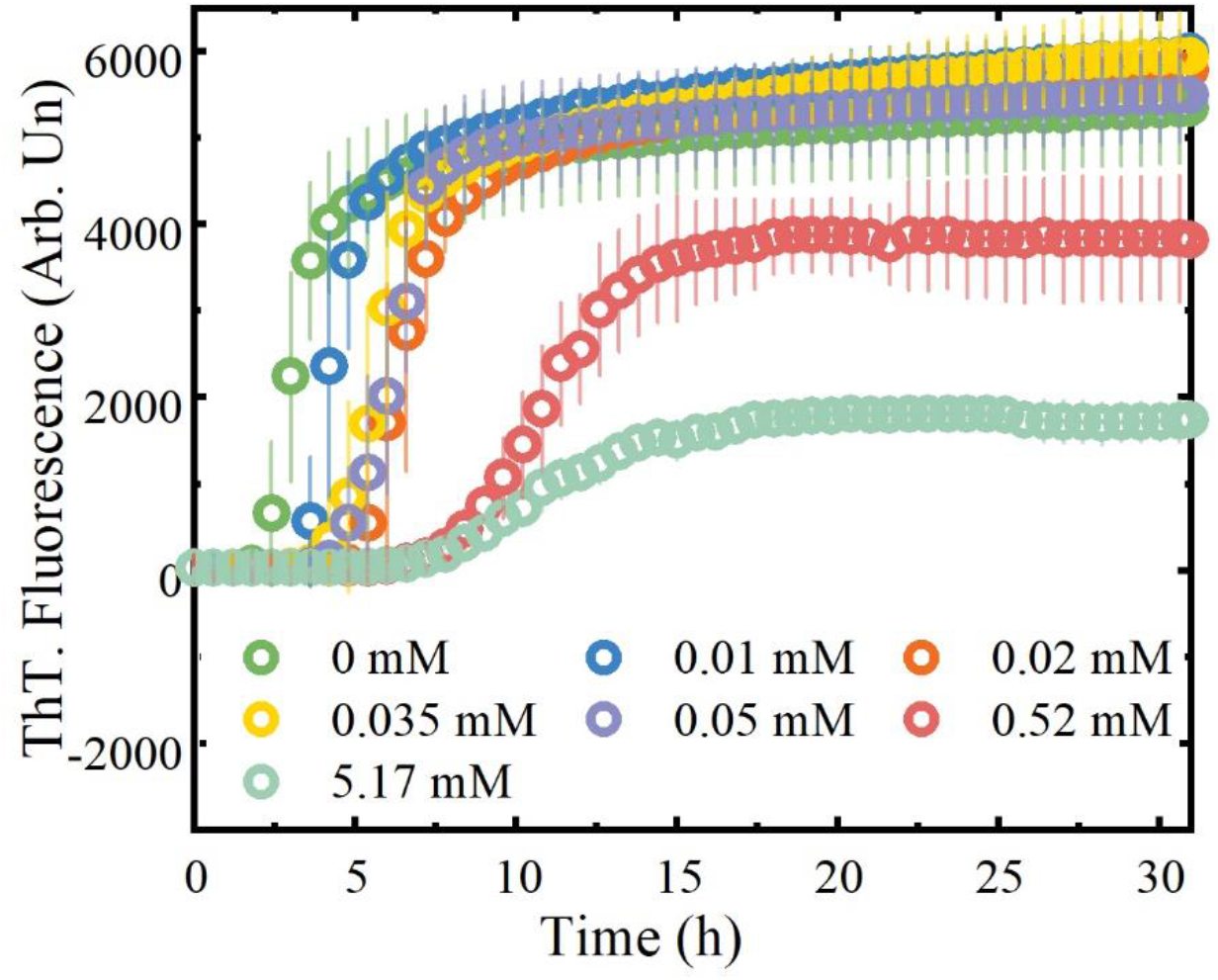
ThT fluorescence kinetics for insulin (3 mg/mL) at different concentrations of PS80 in 20 % (v/v) acetic acid, 0.5M NaCl, pH 1.7 (n=4), incubated at 45 °C for 42 hrs.

### Circular Dichroism (CD) Spectra of Insulin at Increasing PS80 Concentrations Prior to Thermal Incubation

Secondary structure of insulin dissolved in 20 %(v/v) acetic acetic acid, 0.5 M NaCl solution, pH 1.7, at increasing concentration of PS80 was recorded by CD. CD spectra were collected at 20 °C on a Chirascan CD spectrometer (Applied Photophysics, Leatherhead, Surrey, UK), the settings were as follows: 1 nm data interval, 1 nm bandwidth, 2 nm/s scanning speed. 3 mg/mL insulin was dissolved in solutions containing PS80 in the concentration range from 0 mM to 5.17 mM. Cuvettes with a 50 µm path length (Type 19 Demountable U-Shaped Circular Dichroism Cuvette, FireflySci Cuvette Shop, NYC, NY, USA) were used in this experiment. Far-UV CD of insulin was recorded in the range of 190-260 nm. The presented data for each sample are an average of five scans. Figure S5 shows that, apart from a slight variation at 192 nm, the addition of PS80 did not significantly change the overall shape of the CD spectrum from 200 nm to 260 nm, with all spectra mainly showing a dominant α-helix structure prior to incubation.

**Figure S5.**
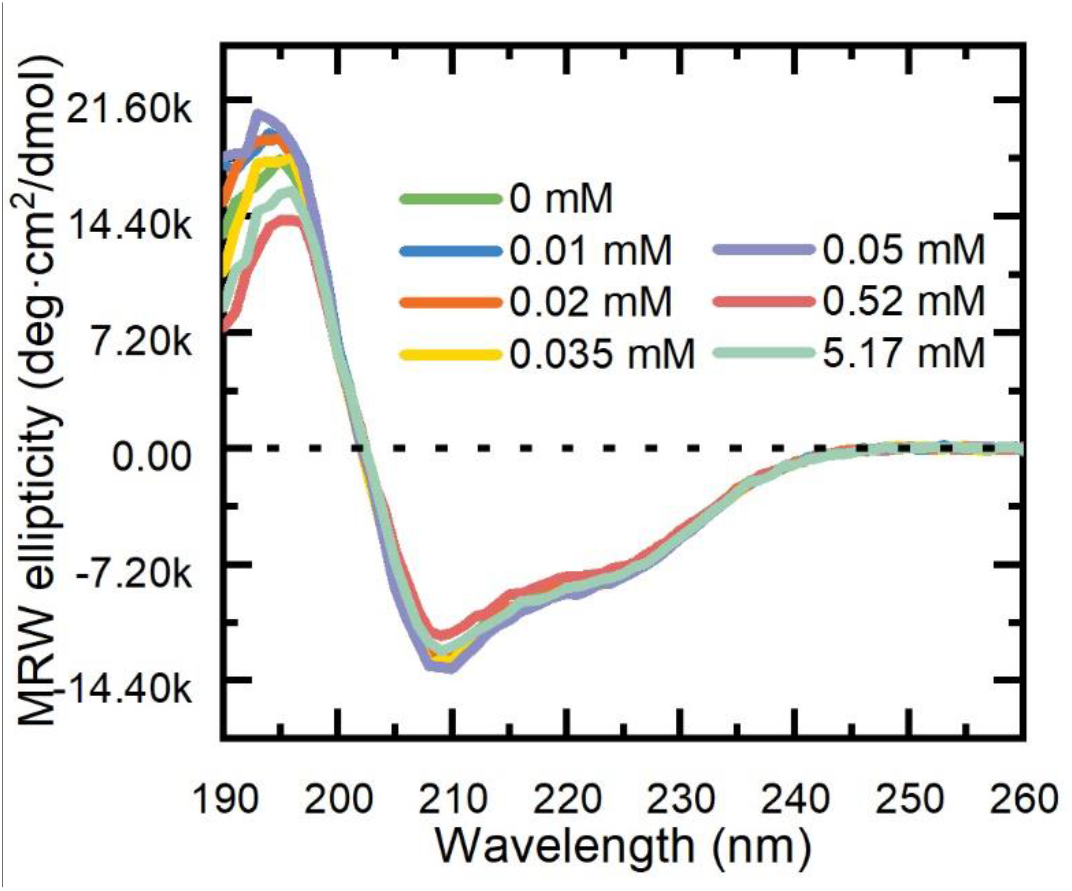
Far UV circular dichroism (CD) spectra of insulin at increasing PS80 concentrations before formation of spherulites. Insulin was dissolved in 20 %(v/v) acetic with 0.5 M NaCl addition, pH 1.7, the concentration of PS80 is shown in the legend in the figure. Spectra are the result of the mean of five scans.

### Qualitative Analysis of ThT Fluorescent Lifetime Bound to Insulin Spherulites Formed at Different PS80 Concentrations

Using phasor analysis each pixel of the intensity images is mapped to a point in the phasor plot corresponding to the measured fluorescence lifetime. Single exponential lifetimes lie on the so-called “universal circle”; long lifetimes are located near the origin (0 on the x axis), while short lifetimes are shifted on the circumference toward the bottom right intersection with the x axis (1 on the x axis) (Digman et al. 2008). Phasors placed inside the universal circle indicate that ThT lifetime in these conditions is characterised by by mutiple exponential decays. In the lifetime maps (Figure S6a) pixels are colored by selecting colored ROIs in the polar plot (Figure S6b). When overlapping the populations (Figure S6c), 3 distributions can be observed: one, in green, corresponds to the samples below CMC (0.05 mM) of PS80 with shorther fluorescence lifetime. A population in red corresponds to the samples above the CMC of PS80 and has the longest lifetime, while an intermediate population in pink corresponds to the samples formed at the CMC value. When fluorescence lifetime distributions of samples at different concentration of PS80 are plotted on the same phasor plot, they lie on a straight line. Hence, a quantitative analysis shown in Figure 5 can be performed splitting the population into two single exponential fluorescence decays (De Luca et al. 2020).

**Figure S6.**
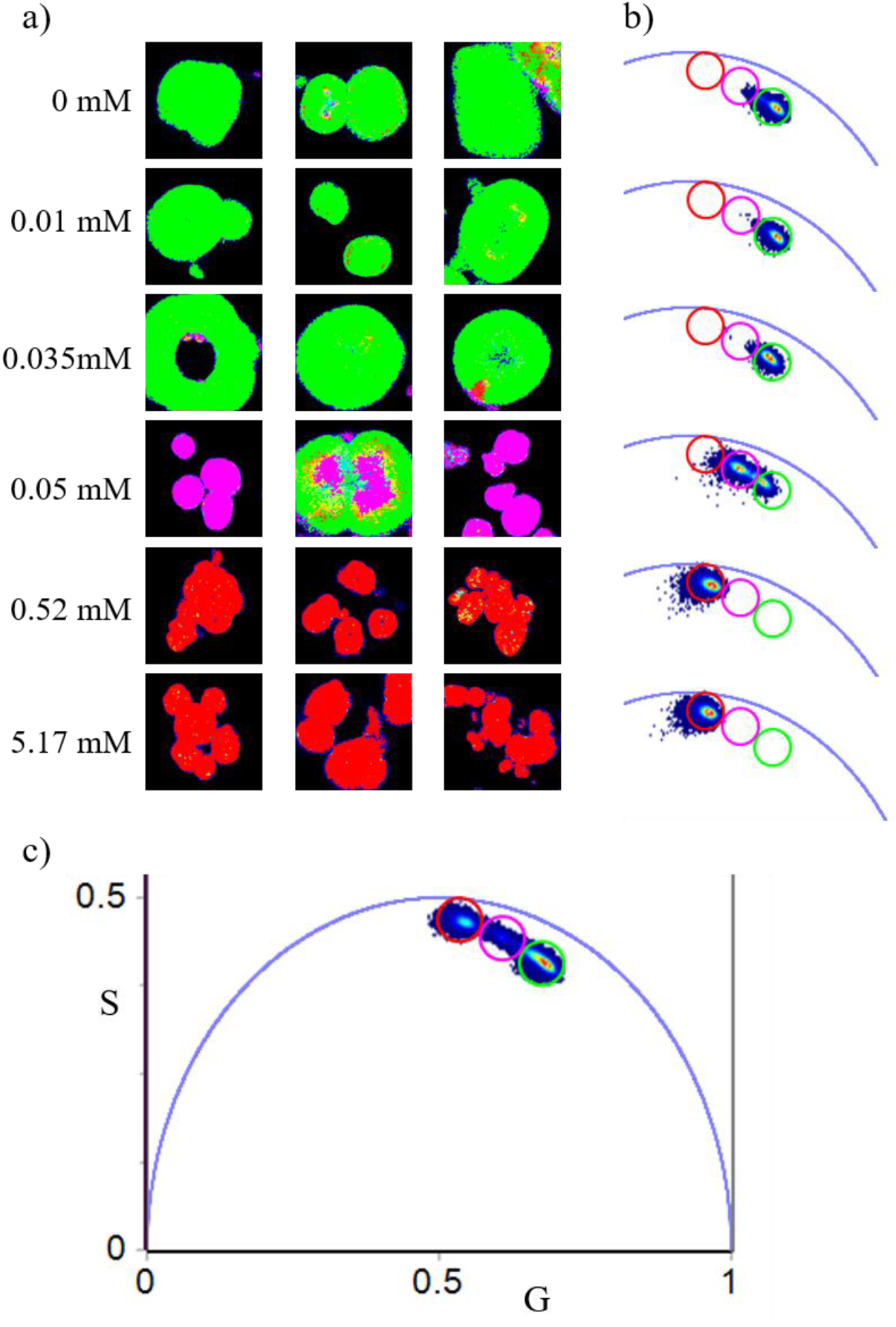
a) 256 × 256 qualitative lifetime maps of spherulites formed at different concentrations of PS80 (0, 0.01,0.035, 0.05, 0.52 and 5.17 mM) and stained with ThT 80 µM where pixels are colored according to the ROIs at the phasor plot. b) Phasor plot of each sample showing the different populations at different PS80 concentrations. c) Superposition of all populations clearly showing that they lie on a straight line.

### Spherulites Dissociation and the Release of Soluble Insulin in Solutions at pHs 1.7, 5 And 7

The tested spherulites did not dissociate in buffers at pH values of 1.7, 5, and 7. The pH 1.7 buffer corresponds to the acetate buffer used in the spherulite preparation process (without PS80). Acetate buffer (pH 5) was prepared by dissolving 288.6 mg sodium acetate (Merck, ≥99.0%, Darmstadt, Germany) and 83.9 µL acetic acid into 50 mL purified water, adjusted to pH 5 using 5 N HCl. The phosphate buffer (pH 7) was prepared by dissolving 410.1 mg disodium hydrogen phosphate (Merck, ≥99.0%, Darmstadt, Germany) and 253.0 mg monosodium dihydrogen phosphate (Merck, ≥ 99.0 %, Saint Louis, MO, USA) in 50 mL purified water, adjusted to pH 7 using 5 N HCl and 5 N NaOH.

**Figure S7.**
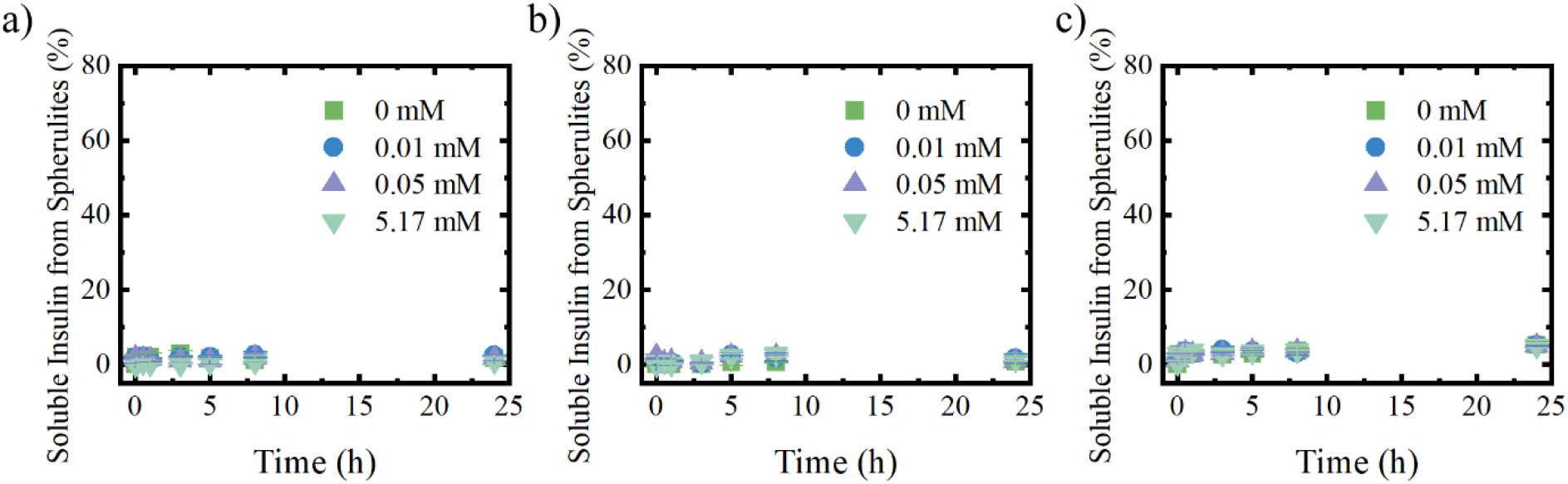
Soluble insulin released from spherulites. Spherulites dispersed in media with different pH values, a) pH 1.7 (20 %(v/v) acetic acid with 0.5 M NaCl), b) pH 5 (acetate buffer, 0.1 M), c) and pH 7 (phosphate buffer, 0.1 M). The amount of insulin released from sphetulites is reported as the percentage of the initial mass of insulin spherulites. Mean ± SD, n=3.

## Notes

### Competing Interest Statement

The authors have declared no competing interest.

### Summary of Updates

We included an extra set of data (Figure 1)and the text has been slightly modified to increase the readability of the manuscript. Supplementary materials are now included together with the main text in a single file.

